# A Common Cause for Nystagmus in Different Congenital Stationary Night Blindness Mouse Models

**DOI:** 10.1101/2023.04.24.538135

**Authors:** Maj-Britt Hölzel, Beerend H. J. Winkelman, Marcus H. C. Howlett, Wouter Kamermans, Chris I. De Zeeuw, Maarten Kamermans

## Abstract

In *Nyx*^*nob*^ mice, a model for congenital nystagmus associated with congenital stationary night blindness (CSNB), synchronous oscillating retinal ganglion cells (RGCs) lead to oscillatory eye movements, i.e., nystagmus. Given the distribution of *mGluR6* and *Cav1*.*4* in the retina as well as their clinical association with CSNB, we hypothesize that *mGluR6*^*-/-*^ and *Cav1*.*4*^*-/-*^ mutants show, like the *Nyx*^*nob*^ mouse, oscillations that originate in the A_II_ amacrine cells (A_II_ ACs). Using eye movement and multi-electrode array (MEA) recordings of RGCs we show that the nystagmus as well as the underlying RGC oscillations are also present in *mGluR6*^*-/-*^ and *Cav1*.*4*^*-/-*^ mice. Yet, we find that the oscillations in the *mGluR6*^*-/-*^ and *Cav1*.*4*^*-/-*^ mutants slightly differ from each other and also from those of the *Nyx*^*nob*^ mice. Moreover, each of the three mutations likely impacts the membrane potential of the A_II_ ACs differently. Together our results indicate that nystagmus and oscillating RGCs are generalizable features associated with CSNB mutations localized at the photoreceptor-bipolar cell synapse.

## Introduction

Patients with congenital stationary night blindness (CSNB) often have congenital nystagmus as well (Pieh *et al*., 2008; Bijveld *et al*., 2013). These involuntary, oscillating eye movements occur shortly after birth and vary from pendular to jerk nystagmus (Optican & Zee, 1984; Pieh *et al*., 2008). Previously, we reported that *Nyx*^*nob*^ mice, a CSNB mouse model (Pardue *et al*., 1998; Gregg *et al*., 2003), have a pendular nystagmus that results from the oscillatory output of retinal ganglion cells (RGCs, Winkelman et al., 2019). This oscillatory firing of RGC is driven by A_II_ amacrine cells (A_II_ ACs), which are most likely depolarized outside of their normal working range due to the lack of ON-bipolar cell (ON-BC) input (Winkelman *et al*., 2019).

The photoreceptor to ON-BC synapse is a highly specialized metabotropic glutamatergic synapse (Fig 1). Depolarization of a photoreceptor opens its Cav1.4 calcium-channels and the resulting influx of Ca^2+^ into its synaptic terminal ultimately leads to glutamate release (Schmitz & Witkovsky, 1997). The released glutamate diffuses across the synaptic cleft and binds to the postsynaptic metabotropic glutamate receptors (mGluR6, Masu et al., 1995). Activation of this receptor triggers a G-protein coupled cascade, which leads to the closure of TRPM1 channels resulting in ON-BC hyperpolarization (Shen *et al*., 2009; Morgans *et al*., 2009). The function of this synapse depends crucially on the close interaction of all the synaptic proteins involved.

**Figure 1:**
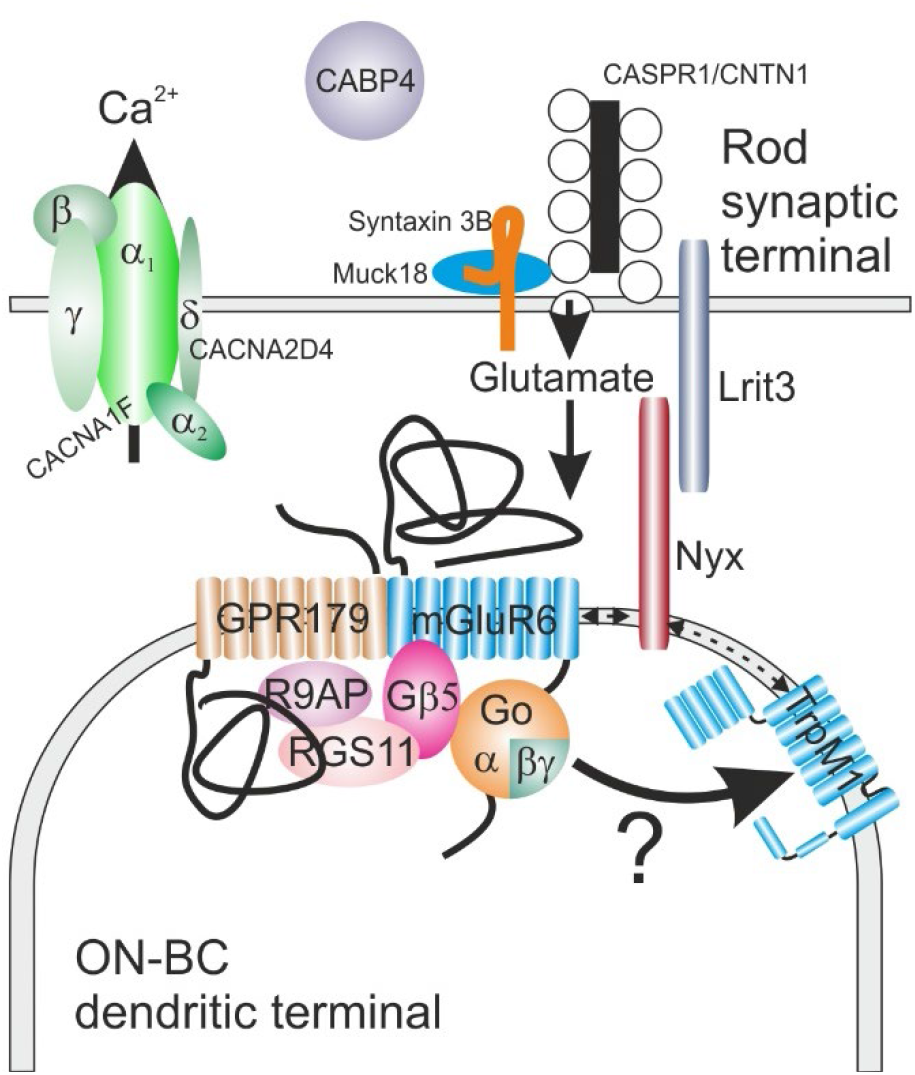
Schematic overview of the photoreceptor to ON-bipolar cell synapse

Mutations of the various proteins involved in this synapse will each affect the ON-BC differently. Post-synaptically, there are varying degrees of interdependency between the elements of the signalling cascade. For instance, elimination of mGluR6 disrupts both the expression levels and subcellular targeting of most signalling molecules so far examined. By contrast, when the scaffolding protein nyctalopin is lost, TRPM1 is no-longer correctly trafficked and localised to the dendritic tips but other elements of the signalling cascade remain unaffected (Pardue *et al*., 1998; Gregg *et al*., 2007, 2014; Cao *et al*., 2009, 2011; Pearring *et al*., 2011; Ray *et al*., 2014; Martemyanov & Sampath, 2017). Furthermore, the post-synaptic receptor complex remains intact for Cav1.4 mutations but the presynaptic signalling is impaired. These different conditions will likely lead to different membrane potentials of the ON-BC and possibly to slightly different phenotypes of CSNB

There are several CSNB mouse models, each with a different photoreceptor to ON-BC synapse specific protein mutated (Zeitz *et al*., 2015). As mutations in each of these different proteins likely affects the ON-BC differently, it is possible that each mutation presents with a phenotypically distinct nystagmus. To test this idea, we studied eye movements and RGC activity in two CSNB mouse models: *Cav1*.*4*^*-/-*^ and *mGluR6*^*-/-*^ mice and compared the results with those of *Nyx*^*nob*^ mice. We found that all CSNB mouse models had a disturbed OKR, nystagmus and oscillating RGCs. However, each mouse model displayed phenotypically distinct features including different eye movements, and RGC oscillation frequencies. Furthermore, we found that flashes of light at different intensities, which should polarize the A_II_ ACs to differing degrees, changed the oscillation frequency of *Nyx*^*nob*^ RGCs.

## Material & Methods

### Animals

All animal experiments were carried out under the responsibility of the ethical committee of the Royal Netherlands Academy of Arts and Sciences (KNAW) acting in accordance with the European Communities Council Directive of 22 September 2010 (2010/63/EU). The experiments were performed under the license numbers AVD-801002016517 and AVD-80100202115698, issued by the Central Committee for Animal Experiments of the Netherlands.

*mGluR6*^*-/-*^ and *Nyx*^*nob*^ mice were obtained from the McCall lab (University of Louisville, Loiusville, USA) and *Cav1*.*4*^*-/-*^ mice from the Koschak lab (University of Innsbruck, Innsbruck, Austria). All mice were either in a C57BL/6JRj background with or without a GFP label on ON-dsRGC coding for upward motion (SPIG1^+^ mice,Yonehara et al., 2009). Since the Nyx mutation is x-linked, only male mice in the age range of 5-71 weeks were used for the experiments. Room lights were timed on a 12/12 h light-dark schedule, and experiments took place during daytime. Mice had *ad libitum* access to food and water.

### Eye movement recordings

#### Surgical preparation

Prior to the start of experiments, adult animals were equipped with a head-fixation pedestal, an anodized aluminium bit with an integrated magnet, attached to the parietal and frontal bones of the skull using dental cement (Super Bond C&B, Sun Medical, Japan). Surgery was performed under general isoflurane/O_2_ and topical anaesthesia (bupivacaine). Analgesia was offered by subcutaneous injection of carprofen (5 mg/kg). The recovery time was at least two days after pedestal surgery. During experiments, the animal was placed head-fixed in the experimental setup using a custom-made adapter, which allowed panoramic vision.

#### Optokinetic stimulus setup

Optokinetic stimuli were displayed on two Benq XL2420t high-performance monitors (120 fps, gamma-1.745) that were placed in V-formation around the animal. The closest distance between the screen surface and the mouse head was 16.5 cm. Screen dimensions were 56.9 × 33.8 cm (combined field of view: 240° × 50°). For measurements in darkness, the displays were switched off. Mickelson contrast of the grayscale grating stimuli was 99.67%. Visual stimuli consisted of sine wave gratings (mean luminance: 71.6 cd/m^2^) and homogeneous grayscale images (mean luminance: 71.6 cd/m^2^).

#### Eye movement recordings

Eye movements were recorded using an infrared video tracking system (JAI RM-6740CL monochrome CCD camera, 200 fps, 850 nm illumination). Pilocarpine (2%) eye drops were used to reduce pupil size in case the pupil was too large to track. Online image analysis extracted the location of pupil edges and corneal light reflections from each frame using custom-built software for LabVIEW (National Instruments, Austin, TX, USA). Eye position was computed from the relative distance between pupil centre and corneal reflections of the infrared LEDs (Stahl *et al*., 2006) and pupil size (Stahl *et al*., 2000; Stahl, 2002). Epochs containing saccades, eye blinks, and motion artefacts were excluded from analysis. Eye velocity was smoothed using a Gaussian smoothing kernel with a SD of 7.5 ms (25-Hz cut-off). All OKR measurements were done monocular, during which, the contralateral eye was covered by a miniature blackout cap.

#### Analysis of eye movement recordings

The power spectral density (PSD) was computed from angular eye velocity using Welch’s method with a 4-s window length, 75% overlap between windows, and a Hann window function. The average PSD for each genotype was computed using the geometric mean (Pintelon *et al*., 1988; Attivissimo *et al*., 2000).

### Multielectrode RGC recordings

#### Retinal dissection

Multielectrode RGC recordings were performed as described previously (Winkelman *et al*., 2019; Hölzel *et al*., 2022). After at least one hour of dark adaptation, mice were sedated using a mixture of CO_2_/O_2_ and ultimately euthanized by cervical dislocation. All procedures were carried out under dim red light. The eyes were extracted from the eye socket and placed in room temperature Ames’ medium (Sigma-Aldrich, St Louis, MO). Next, the cornea and lens were removed by making an insertion around the *ora serrata* using fine spring scissors. As much vitreous humour as possible as well as the sclera were removed using fine forceps. Four small insertions were made and the retina was flat mounted on a filter paper annulus (1 mm inner radius; 0.8 µm hydrophilic MCE MF-MilliporeTM membrane filter, Merck Millipore Ltd., Tullagreen, Ireland). The retina was then mounted photoreceptor cell side up on a perforated 60-electrode MEA chip (60pMEA200/30iR-Ti, Multichannel systems, Reutlingen, Germany) in a recording chamber mounted on a Nikon Optiphot-2 upright microscope and viewed under IR with an Olympus 2x objective and video camera (Abus TVCC 20530). During the experiment the retina was continuously superfused with Ames’ medium gassed with a mixture of O_2_ and CO_2_ at a pH of 7.4 and a temperature of 29-36°C. Before the recording started, an acclimatisation period in the dark for a duration of 15 minutes was given to ensure stable recordings.

#### Data acquisition

The extracellular RGC activity was acquired using MC rack (Multichannel systems, Reutlingen, Germany) at a sampling frequency of 25 kHz. The data was then zero-phase bandpass filtered (250-6250 Hz) with a fourth-order Butterworth filter in Matlab (Mathworks, Natick, MA, USA). Subsequently the spiking activity was sorted manually into single unit activity using the Plexon offline sorter (Plexon, Dallas, TX, USA) based on the first two principal components versus time. For the extraction of spikes from the background, a spike detection amplitude threshold of > 4σ_n_ was used as criteria. Hereby σ_n_ is defined as an estimation of the background noise and x being the bandpass-filtered signal (Quiroga *et al*., 2004).

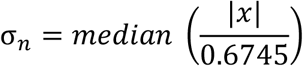

#### Optical stimulator

Full field white light flashes were generated using Psychophysics Toolbox Version 3 (Brainard, 1997) and presented to the photoreceptor side of the retina with a custom-modified DLP projector (for details of modification see Winkelman et al., 2019; Light Crafter 4500, Wintech, Carlsbad, CA, USA). For WT, *Nyx*^*nob*^ and *Cav1*.*4*^*-/-*^ retinas, the full field light flash had a duration of 500 ms. Since the oscillations in mGluR6^*-/-*^ mice were much slower than in the other mouse models the flash duration was extended to 1000 ms. In all cases the light flash was preceded by a 500-and followed by a 1000 ms period of darkness and the stimulus was repeated 100 times. White light stimuli consisted of equal quantal output of the red (625 nm), green (530 nm) and blue (455 nm) LEDs. The maximal light intensity was 8.60 10^19^ quanta m^-2^ s^-1^. For the study of the effect of different light intensities on the *Nyx*^*nob*^ RGC oscillation frequency, two light intensities were used: 9.02 10^18^ quanta m^-2^ s^-1^ and 8.60 10^19^ quanta m^-2^ s^-1^.

#### Data analysis

To identify spontaneous oscillatory RGCs activity, the firing activity in the dark was recorded a period of at least 15 minutes. For each cell the activity was binned in 1 ms bins, and then divided into 5 s non-overlapping epochs. Each epoch was baseline subtracted and the autocorrelation and PSD determined. Based on this data the average autocorrelation and Welch’s PSD was calculated for each cell. The RGCs of one the five *Cav1*.*4*^*-/-*^ retinas oscillated at much lower frequencies than the other four and was therefore excluded from the analysis. The mean light responses were calculated over the 100 stimulus repetition for each cell. 500 ms of the mean light response of each cell that occurred after 50 ms after the light onset was zero padded to 2 s and used to calculate the Welch’s PSD.

The median oscillation frequency of the eyes was plotted against the median oscillation frequency of the RGCs and fitted with a linear regression model.

#### Spike train synchrony

Synchronisation between spike trains were quantified using SPIKE-Synchronization. This time-scale independent and parameter-free metric uses Inter-Spike-Interval derived coincidence windows to generate a similarity scores. A similarity score of 1 only occurs if each spike in a train has a matching spike in the other train(s) whereas a score of 0 only occurs if the spike trains do not contain any coincidences (for more details see: Kreuz et al., 2015; Satuvuori et al., 2017). The analysis routine was used within cSPIKE, a publicly available Matlab-based spike train analysis software package (https://thomas-kreuz.complexworld.net/source-codes/cspike).

We generated two distributions of similarity scores for each retina. The first distribution was of similarity scores between spike trains of spontaneous activity occurring in the dark for units within the same retina. The spontaneous activity was divided into 5s long non-overlapping periods and for each of these periods the similarity scores for each unit vs every other unit determined (e.g. a 15 min recording of 30 units resulted in 30*30*180 similarity scores). For these same units we also generated a second distributions of similarity scores, this time comparing their spike trains with those of units from different retinas in the other treatment groups. For example, units from a single *Cav1*.*4*^*-/-*^ retina were compared with unrelated units from all the *Nyx*^*nob*^, WT and *mGluR6*^*-/-*^ retinas, again using 5s long non-overlapping periods. This second population of similarity scores was used to estimate the chance distribution of similarity scores occurring for units whose activity is unrelated.

For each retina, we quantified the distance between their two distributions of similarity scores using the Hellinger Distance. The relative probability distribution of the similarity scores were distributed over 51 equally spaced bins ranging from 0 to 1 in 0.02 increments, then for the two discrete probability distributions P =(p_1_,….p_k_) and Q=(q_1_,….q_k_), their Hellinger Distance defined as

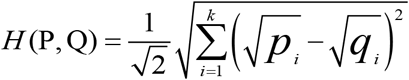

where 1 indicates there is no overlap of the populations and 0 indicates the two populations fully overlap.

#### Statistical Analysis

Statistical analysis was performed using SPSS Statistics 28 (IBM Corp., Armonk, NY, USA) and Origin Pro 8 (Northampton, MA, USA). Since the distribution of the eye movement oscillations frequencies violated the assumption of normality (Shapiro Wilk), statistical differences were assessed using an Independent-Samples Mann-Whitney U test. All values are given as median ± MAD and a statistically significant difference was assumed with a p-value < 0.05. Statistical differences in the Hellinger distances were tested by a one-way ANOVA with a posthoc Bonferroni test. To test whether there is a statistical significant effect of the light intensity on the oscillation frequency we used a Wilcoxon signed-rank test.

## Results

### Oscillating eye movements in *mGluR6*^*-/-*^ and *Cav1*.*4*^*-/-*^ mice as compared to those in *Nyx*^*nob*^ mice

To investigate whether there is a common cause for nystagmus in CSNB we first measured the eye movements of WT, *mGluR6*^*-/-*^, *Nyx*^*nob*^ and *Cav1*.*4*^*-/-*^ mice. Fig. 2 shows example raw traces of the horizontal eye position for each of these mice in response to a monocularly presented horizontal moving vertical grating. WT mice show a normal optokinetic response (OKR). They successfully follow the grating in the temporal to nasal direction (6 s to 10 s), keep their eyes still when the stimulus is stationary (4 s to 6 s) and do not follow the grating when it moves in the nasal to temporal direction (0 s to 4 s).

**Figure 2:**
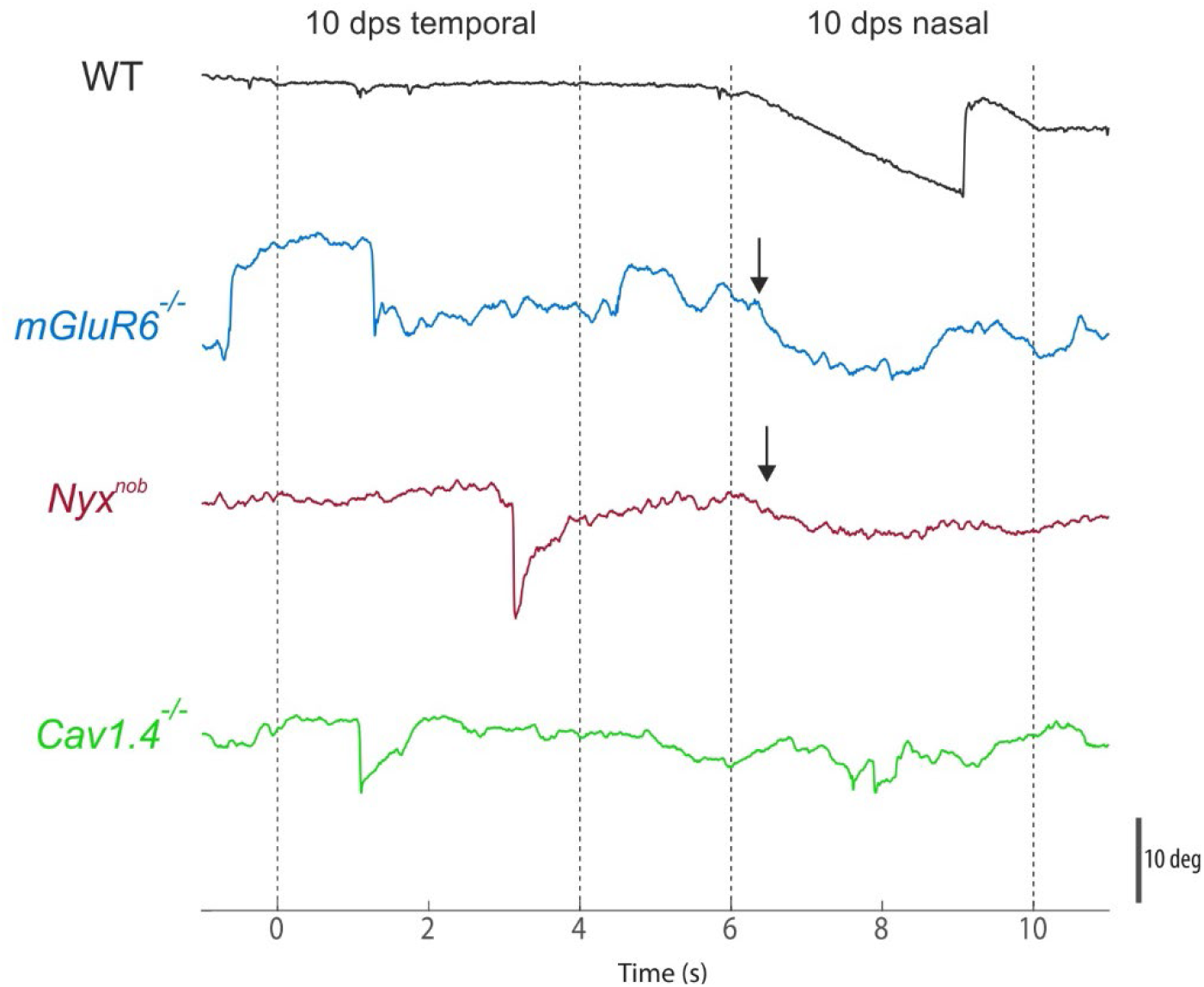
Example horizontal eye movement traces for a WT, *mGluR6*^*-/-*^, *Nyx*^*nob*^ and *Cav1*.*4*^*-/-*^ mouse in response to a vertical sinusoidal grating presented to the left eye. All three mouse models show small oscillatory eye movements.

On the other hand, the OKRs of *mGluR6*^*-/-*^, *Nyx*^*nob*^ and *Cav1*.*4*^*-/-*^ mice were severely impaired and had had different characteristics in each mutant. *mGluR6*^*-/-*^ and *Nyx*^*nob*^ mice initially follow the gratings moving in the temporal to nasal direction (arrows), but were unable to maintain this eye movement (also see Winkelman et al., 2019). Furthermore, their eyes tended to drift when the grating stopped moving. The *Cav1*.*4*^*-/-*^ mice lacked an OKR altogether but did exhibit a random slow drift in the eye position. However, all mutants showed the presence of small-amplitude high-frequency eye movement oscillations i.e. pendular nystagmus. In this study we will focus only on these small-amplitude eye-movement oscillations and not discuss the OKR further.

Using PSD plots, we quantified the nystagmus under four stimulus conditions: darkness, homogeneous illumination, vertical gratings moving horizontally and horizontal gratings moving horizontally (Fig. 3). While the eyes of *mGluR6*^*-/-*^ and *Nyx*^*nob*^ mice only oscillate when stimulated with a vertical grating, *Cav1*.*4*^*-/-*^ mice show oscillatory eye movements under all stimulus conditions (arrows). Furthermore, the frequency of the eye movement oscillations during the vertical grating condition differs between the three genotypes (median ± MAD; *mGluR6*^*-/-*^: 3.4 ± 0.38 Hz (N=12); *Nyx*^*nob*^: 5.0 ± 0.25 Hz (N=7); *Cav1*.*4*^*-/-*^: 8.0 ± 0.38 Hz (N=6); Mann-Whitney U, p_mGluR6,Nyxnob_= 0.001; p_mGluR6,Cav1.4_= 1.8*10_-4_; p_Nyxnob,Cav1.4_= 0.001). Additionally, the power spectra of all knockout mice also show a low frequency peak representing a slow eye movement drift, which will not be discussed further. WT mice on the other hand do not exhibit comparable eye movement oscillations although a very small peak at around 5 Hz may be observed sometimes in their power spectrum.

**Figure 3:**
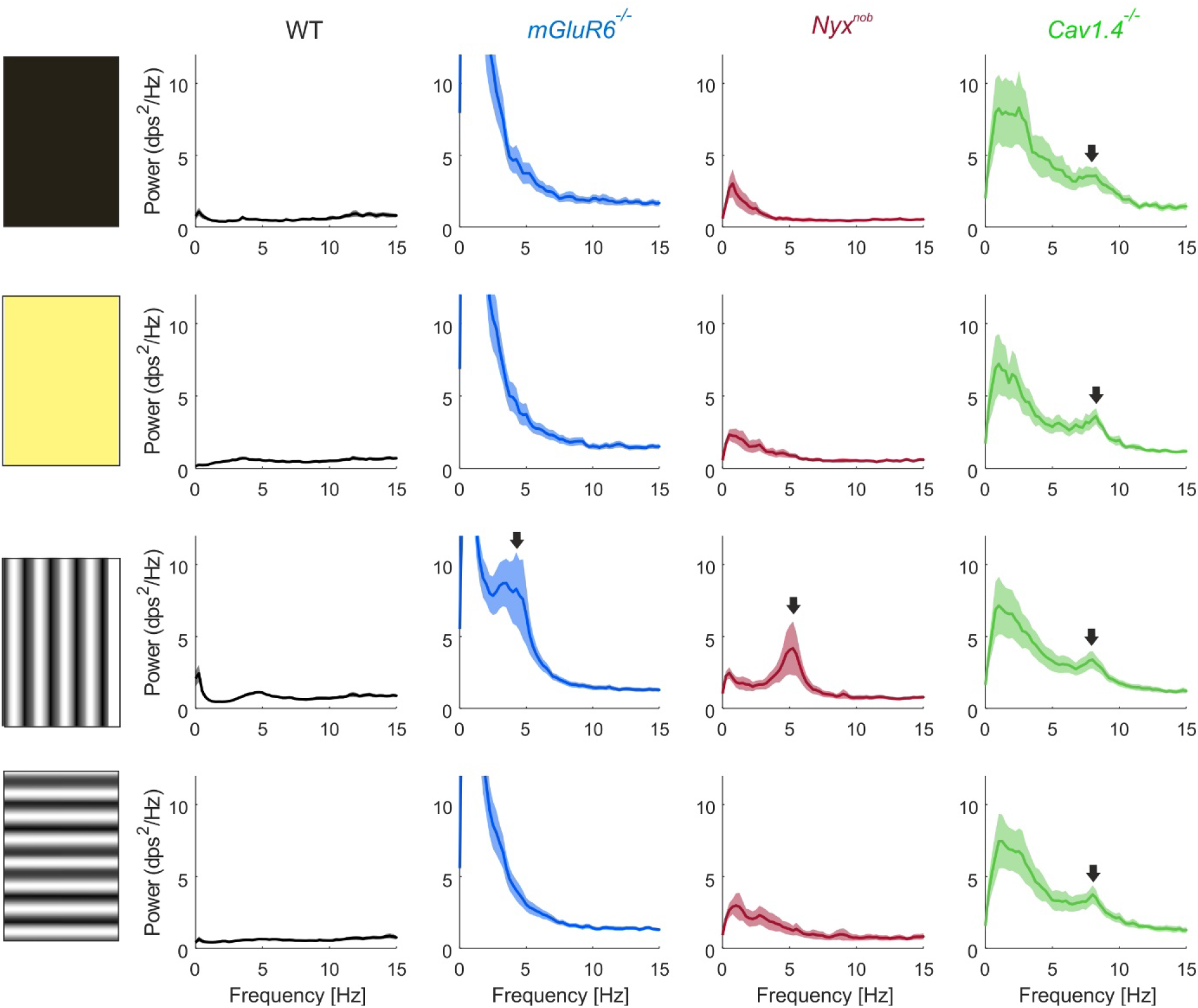
Power spectra comparing the horizontal eye movements of WT (n=7), *mGluR6*^*-/-*^ (n=12), *Nyx*^*nob*^ (n=7) and *Cav1*.*4*^*-/-*^ (n=7) mice in darkness, under homogenous light as well as with vertical and horizontal bar stimulations. Only stimulation with a vertical grating induces oscillatory eye movements in *mGluR6*^*-/-*^ and *Nyx*^*nob*^ mice while the eyes of *Cav1*.*4*^*-/-*^ mice are oscillating under all stimulus conditions.

Taken together, the eye movement data shown in Fig. 3 reveals phenotypical differences between the various CSNB mutants. How does this correlate with the RGC activity?

### Spontaneous oscillation frequencies of *mGluR6*^*-/-*^, *Cav1*.*4*^*-/-*^ and *Nyx*^*nob*^ mice differ

Previously, we have shown that the RGCs of *Nyx*^*nob*^ mice fire in an oscillatory fashion with a frequency closely matching that of their pendular nystagmus. Causality of this relation was shown using retinal specific pharmacological treatments that changed the oscillation frequency of RGC firing resulting in a similar change in frequency of the nystagmus (Winkelman *et al*., 2019). To test whether a similar relation between RGC oscillations and nystagmus exists in the various mutants, we determined their RGC oscillatory firing frequency.

In the dark, the spontaneous activity (Fig 4A) of many *Nyx*^*nob*^ and *Cav1*.*4*^*-/-*^ RGCs was oscillatory as indicated by the periodic variations in their autocorrelations (Fig 4B) and peaks in their PSD plots (Fig 4C). For *Nyx*^*nob*^ mice, their RGCs oscillated at a lower frequency (7.8 ± 0.25 Hz, median ± MAD; n= 368 RGCs, N=8 mice) compared to those of the *Cav1*.*4*^*-/-*^ mice (11.8 ± 1.25 Hz, median ± MAD; n=135 RGCs, N= 4 mice, Mann-Whitney U, p_Nyxnob, Cav1.4_= 0.000). Unexpectedly, the autocorrelations and PSD of *mGluR6*^*-/-*^ RGCs showed no evidence of oscillatory activity and were instead comparable to that of WT animals.

**Figure 4:**
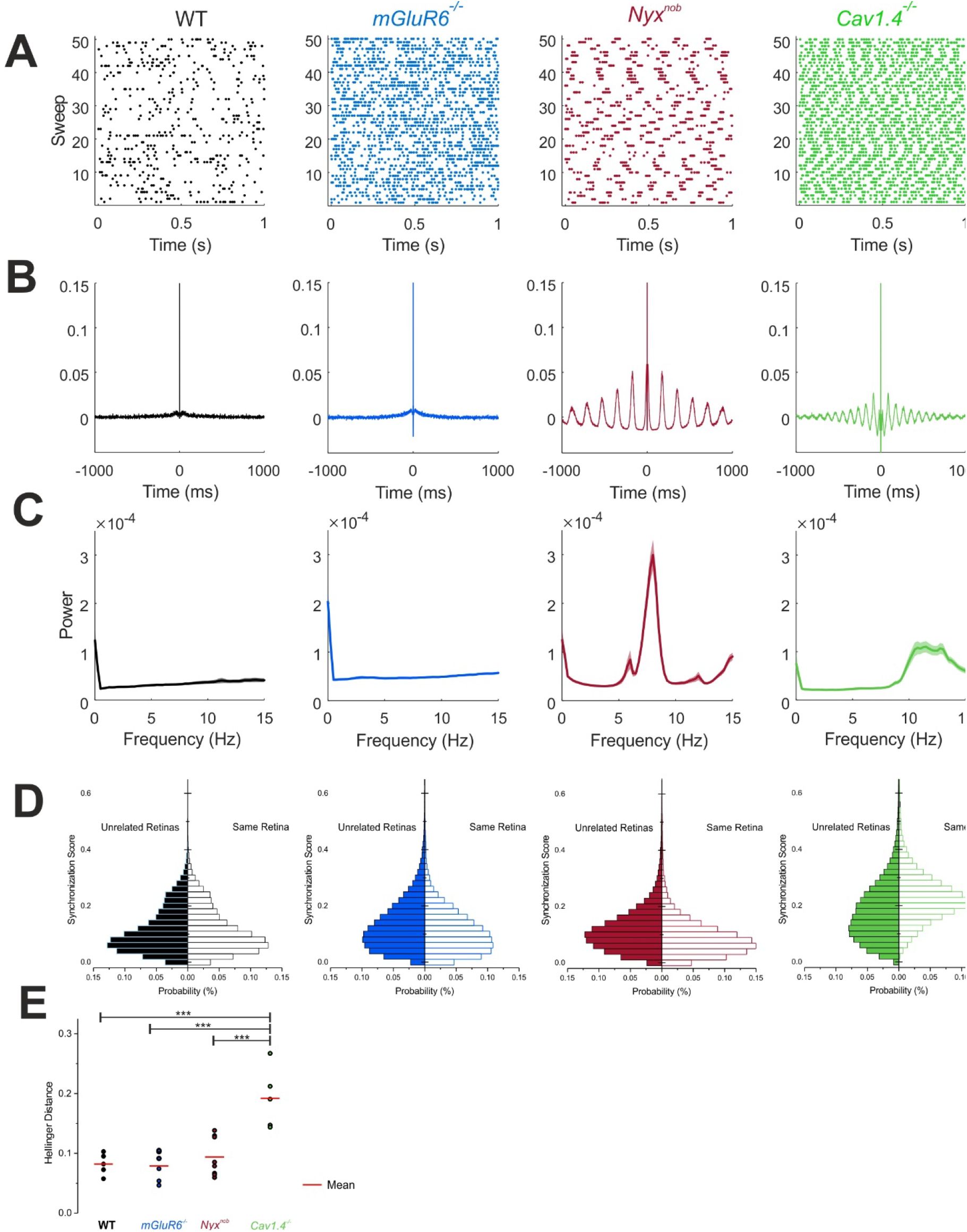
Comparison of the spontaneous RGCs activity of WT, *mGluR6*^*-/-*^, *Nyx*^*nob*^ and *Cav1*.*4*^*-/-*^ mice. (A) Raster plots comparing the extracellularly recorded spontaneous spiking activity for in each case one RGCs from WT, *mGluR6*^*-/-*^, *Nyx*^*nob*^ (red) and *Cav1*.*4*^*-/-*^. (B) Autocorrelation of activity of the same RGC as in (A) showing oscillatory behaviour visible as periodic variations, only for the *Nyx*^*nob*^ and *Cav1*.*4*^*-/-*^ mice but not WT and *mGluR6*^*-/-*^ RGCs. (C) Mean power spectra of the spontaneous activity in WT (n=209 RGCs from N= 5 mice), *mGluR6*^*-/-*^ (n=543 RGCs from N=11 mice), *Nyx*^*nob*^ (n=367 RGCs from N=8 mice) and *Cav1*.*4*^*-/-*^ (n=135 RGCs from N= 4 mice). (D) Comparison of similarity scores as measure for synchrony for spike trains from RGCs from the same versus from another retina. In the *Cav1*.*4*^*-/-*^ retina RGCs are synchronized in the dark which is not the case for the other three mice. (E) Comparison of the Hellinger distance of WT and the three mouse models. ^***^ p < 0.001.

### Light induced oscillations in *mGluR6*^*-/-*^ RGCs

As we showed previously, in *Nyx*^*nob*^ mice, individual RGCs oscillate at similar frequencies, but their activity is asynchronous in the dark. Light stimulation phase-resets the RGC oscillations, and then this synchronized oscillatory retinal output induces oscillating eye movements (Winkelman *et al*., 2019). Could it be, for *mGluR6*^*-/-*^ mice, that light stimulation still induces synchronized RGC oscillations even though their activity is not spontaneously oscillatory in the dark? To answer this question, we studied the responses of RGCs to a light flash (Fig. 5).

**Figure 5:**
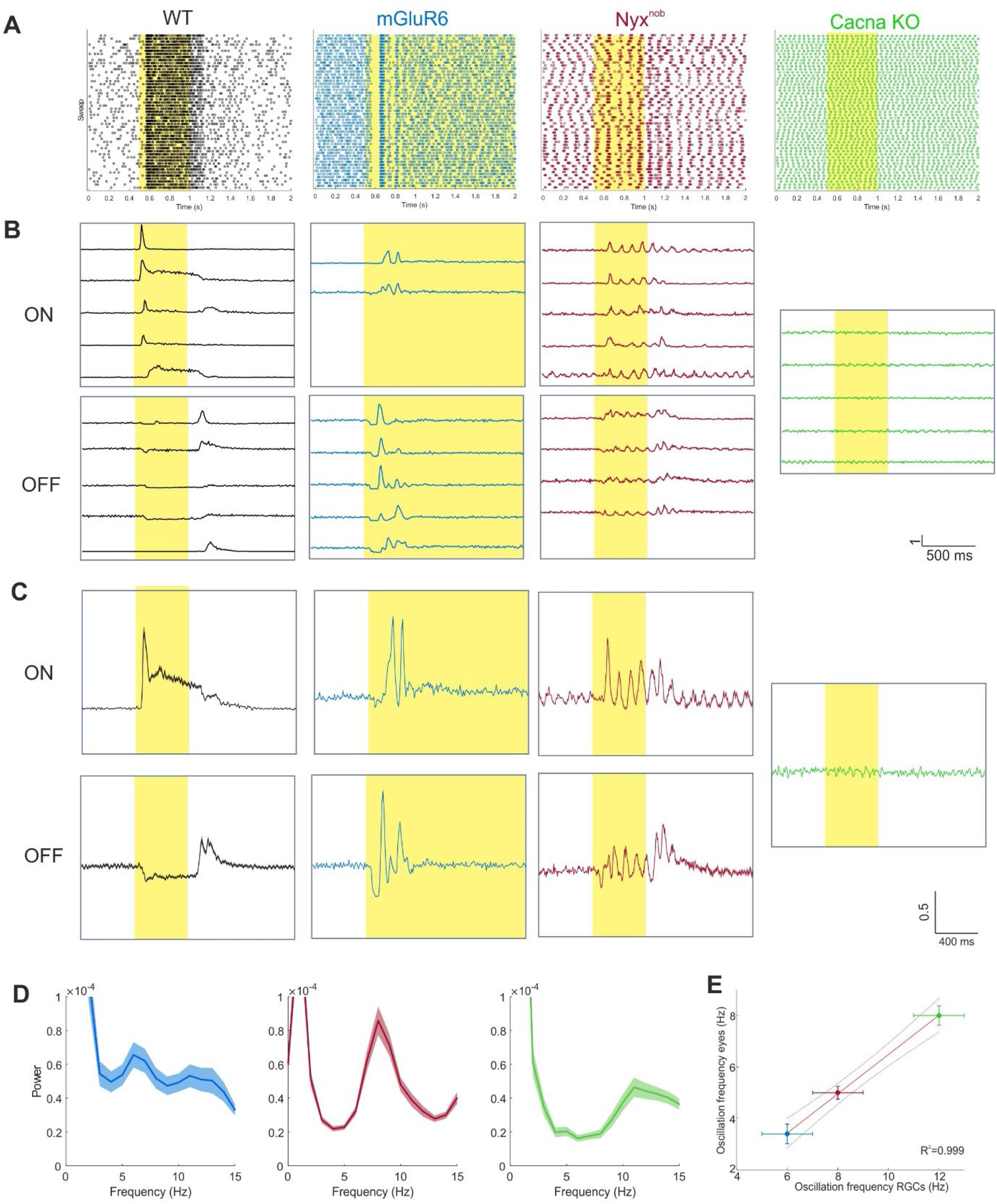
Comparison of the light responses of WT, *mGluR6*^*-/-*^, *Nyx*^*nob*^ and *Cav1*.*4*^*-/-*^RGCs. (A) Raster plots comparing the light responses of in each case one RGCs from WT (black), *mGluR6*^*-/-*^ (blue), *Nyx*^*nob*^ (red) and *Cav1*.*4*^*-/-*^ (green). (B) Example light response traces showing the oscillatory activity of the four genotypes. The *Nyx*^*nob*^ data was replotted from (Winkelman *et al*., 2019). (C) Mean activity during a 500 ms/ 1 s light flash for the RGCs shown in (B). (D) Mean powerspectra of *mGluR6*^*-/-*^ (n=17, N=3), *Nyx*^*nob*^ (n=245, N=7), *Cav1*.*4*^*-/-*^ (n=95, N=4) RGCs. (E) Relation between the median RGC and eye movement oscillation frequencies for the three mouse models. The linear regression model fitted is y = -1.16 + 0.76x.

As expected, a light flash stimulus induced typical ON- and OFF light responses in WT mice without RGCs oscillations (Fig 5A-C). This was not the case for *mGluR6*^*-/-*^ and *Nyx*^*nob*^ mice, which lack the ON light response RGC but showed oscillatory activity briefly after the light stimulation onset. In the *mGluR6*^*-/-*^ mouse only 6.5 % of the RGCs oscillated. As is also the case for *Nyx*^*nob*^ mice, RGCs in the *mGluR6*^*-/-*^ mice showed oscillations directly after stimulus onset. Two groups of RGCs were found that oscillated in counterphase presumably originating from ON- and OFF RGCs (Fig 5 B & C). For *mGluR6*^*-/-*^ mice, the duration of the light flash induced RGC oscillations was much shorter than it was in *Nyx*^*nob*^ mice (Fig. 5 B & C). Furthermore, *Nyx*^*nob*^ RGCs oscillated at a higher frequency than *mGluR6*^*-/-*^ RGCs (median ± MAD, *Nyx*^*nob*^: 8 ± 1.0 Hz, n=245 N=7; *mGluR6*^*-/-*^: 6 ± 1.0 Hz, n=17, N=3, Mann-Whitney U, p_Nyxnob, mGluR6_= 4.3*10^−8^). Hence, light stimulation can induce synchronized RGC oscillations in *mGluR6*^*-/-*^ mice, but they are phenotypically distinct to those of *Nyx*^*nob*^ mice.

### RGCs and eye movements oscillated at similar frequencies

Next, we determined how well the oscillation frequency of RGC activity matched that of the eye movements for the *mGluR6*^*-/-*^, *Nyx*^*nob*^ and *Cav1*.*4*^*-/-*^ animals. For this we used the oscillation frequency of RGC activity during the light flash, and of eye movements during the vertical grating, as these were the only conditions in which oscillatory responses occurred in each of the three models. Fig. 5E shows the relation between the eye movement oscillation frequency and the RGC oscillation frequency. In general, the RGC oscillation frequency was linearly related to the eye movement oscillation frequency, but the RGC oscillation frequency was slightly higher than that of the eye movements.

### RGC oscillations in the *Cav1*.*4*^*-/-*^ are synchronized under all stimulus conditions

*Cav1*.*4*^*-/-*^ mice exhibited nystagmus in the four stimulus conditions (Fig 3), whereas *mGluR6* and *Nyx*^*nob*^ mice show this behaviour only for the vertical grating. This suggests the *Cav1*.*4*^*-/-*^ oscillatory eye movements occur independently from a visual stimulus. The *Cav1*.*4*^*-/-*^ mice also lacked an OKR response (Fig. 2) and their RGCs did not respond to a light flash stimulus (Fig 5A-C), which suggests they are blind. How then can the RGCs oscillations in *Cav1*.*4*^*-/-*^ mice be synchronized? A possibility is, that unlike *mGluR6*^*-/-*^ and *Nyx*^*nob*^ mice, the RGCs in *Cav1*.*4*^*-/-*^ mice display a degree of inherent synchronisation independent of the stimulus condition.

To test this, we assessed to what extent the spontaneous activity of RGCs was synchronised in the dark for each of the mouse models. Using the SPIKE-synchronization metric (https://thomas-kreuz.complexworld.net/source-codes/cspike), we determined two synchronisation-score distributions for each retina (Fig 4D). The first set of scores measured the synchronisation occurring between activity of RGCs from the same retina. The second set was generated by comparing the activity of these same RGCs with that of RGCs from the three other mouse models, giving an estimate of the chance distribution of synchronisation scores for the retina. We then quantified the difference between these two distributions using the Hellinger distance (Fig 4E).

*Cav1*.*4*^*-/-*^ retinas had distributions that shifted towards higher synchronisation scores for RGCs in the same retina versus unrelated retinas, compared to the three other mouse models (Fig 4D, One Way ANOVA, p_Nyxnob, Cav1.4_= 0.0003, p_WT, Cav1.4_= 0.0003, p_Nyxnob, Cav1.4_= 1, p _mGluR6, Cav1.4_= 0.0001, p_mGluR6, Nyxnob_ = 1, p_mGluR6,WT_= 1). Indeed, the lowest Hellinger distance for the *Cav1*.*4*^*-*^ _*/-*_ retinas was greater than the highest Hellinger distance of any other group (Fig 4E).

Furthermore, only the *Cav1*.*4*^*-/-*^ group demonstrated elevated synchronisation scores within the same retina. These outcomes indicate that there is a degree of inherent synchronisation in the spontaneous activity of *Cav1*.*4*^*-/-*^ RGCs that is not present in *mGluR6*^*-/-*^, *Nyx*^*nob*^ or WT conditions.

### A_II_ ACs as potential cause for the difference in eye movement oscillations

Previously, we have shown evidence suggesting A_II_ ACs are driving the oscillatory responses of RGCs (Winkelman *et al*., 2019). A_II_ ACs oscillate when they are outside their normal membrane potential range and the oscillation frequency increases with depolarisation (Choi *et al*., 2014). If this is the case then depolarizing *Nyx*^*nob*^ A_II_ ACs further, should lead to an increase of the oscillation frequency of their RGCs. To test this hypothesis, we recorded the light-induced oscillations of *Nyx*^*nob*^ RGCs at two different light intensities (Fig. 6). Consistent with our expectations, we found that when A_II_ amacrine cells are depolarized more strongly by a light flash of higher light intensity, the RGC oscillation frequency increases by about 1.1 ± 0.91 Hz (median ± MAD, n= 92 RGCs, N= 4 mice). This increase in oscillations frequency was statistically significant (Wilcoxon signed-rank test: Z = -4.179, p= 2.9 *10^−5^, n= 92 RGCs, N= 4 mice).

**Figure 6:**
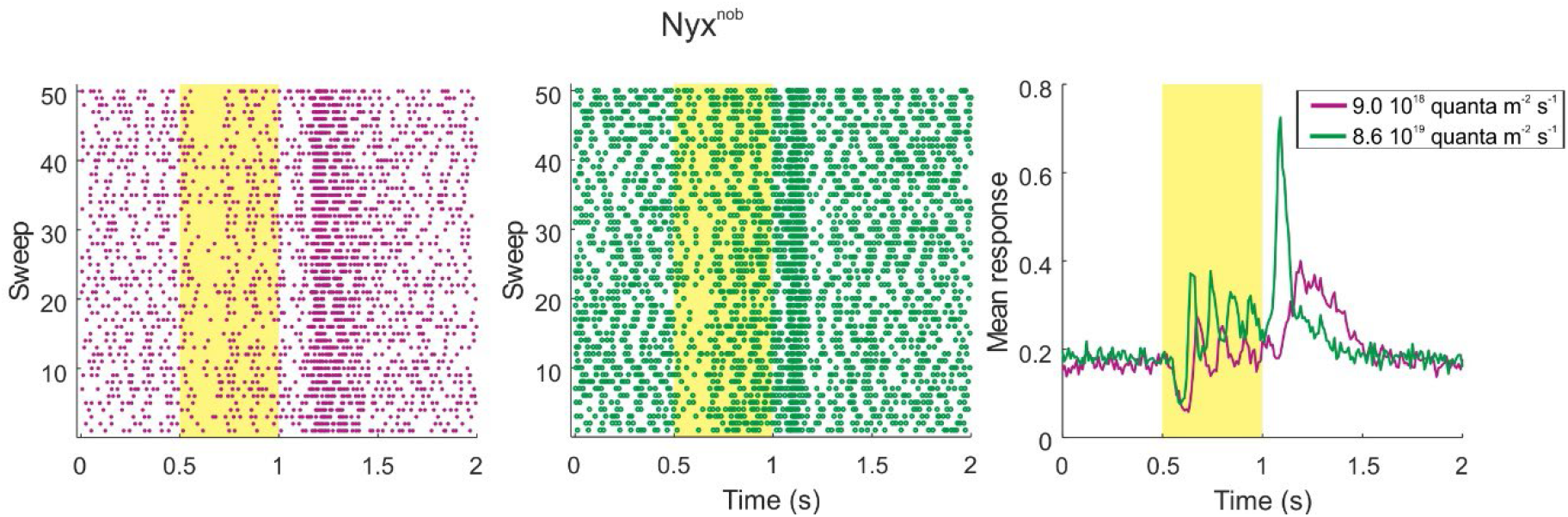
The RGCs oscillation frequency depends light intensity. Left: Raster plots of the light response of one example *Nyx*^*nob*^ RGC for two different light intensities. Right: Mean light response of this example cell over the 100 traces showing that the oscillation frequency increases with higher light intensity.

## Discussion

In the current study, we show that all three mouse models with mutations associated with CSNB, i.e., *mGluR6*^*-/-*^, *Nyx*^*nob*^ and *Cav1*.*4*^*-/-*^ mice, have oscillatory RGCs and eye movements. The oscillation frequencies of the RGCs and eyes movement showed a strong positive linear relationship, while each genotype displayed several phenotypic characteristics. Firstly, the oscillation frequencies associated with each genotype differed, with the lowest frequencies present in *mGluR6*^*-/-*^ mice and highest in *Cav1*.*4*^*-/-*^ mice. Secondly, only vertical gratings induced eye movement oscillations in *mGluR6*^*-/-*^ and *Nyx*^*nob*^ mice, whereas for *Cav1*.*4*^*-/-*^ mice they occurred in all stimulus conditions. Thirdly, in the dark the spontaneous firing of RGCs in *Nyx*^*nob*^ and *Cav1*.*4*^*-/-*^ mice oscillated, but didn’t for *mGluR6*^*-/-*^ mice. Fourthly, light-flash stimuli lead to synchronised RGC oscillations in *Nyx*^*nob*^ and *mGluR6*^*-/-*^ mice, but not in *Cav1*.*4*^*-*^ _*/-*_ mice. And finally, in the dark *Cav1*.*4*^*-/-*^ RGC firing displayed a degree of inherent synchronisation that was not present for *Nyx*^*nob*^ and *mGluR6*^*-/-*^. Below we propose how these different outcomes can arise from a single retinal source.

### Similar mechanism underlying the nystagmus in the three mouse models

Can our previously proposed retinal origin of nystagmus’s model account of the present results (Winkelman *et al*., 2019)? In short, the proposed mechanism works as follows. In *Nyx*^*nob*^ mice A_II_ ACs are depolarized outside their normal working range, which makes them oscillate spontaneously. However, due to small input differences in the various A_II_ ACs, they all oscillate with slightly different frequencies and phases. These oscillations are transmitted to the RGCs, including the ON-DSRGCs. In the dark, the signals from the ON-DSRGCs to the AOS are asynchronous and so the convergent signal generated is not large enough to induce an eye-movement. However, as soon as a light stimulus occurs, the various A_II_ ACs are phase reset, which synchronizes their oscillations and subsequently those of the RGCs as well. Now when the AOS integrates the synchronized RGC inputs, the resulting signal is large enough to induce an eye movement. In turn, this eye movement induces a new visual stimulus, which again synchronizes the retinal oscillations and the loop repeats.

### What causes the differences between the phenotypes of the three mouse models?

Can our proposal account for the distinct phenotypes each of the three CSNB mouse models displayed? These distinct features include: 1) different oscillation frequency of each CSNB model, 2) the absence of spontaneous RGC oscillations in the dark for the *mGluR6*^*-/-*^ mice and 3) the stimulus independent synchronized oscillations of the RGCs in *Cav1*.*4*^*-/-*^ mice?

#### 1) Differences in oscillation frequencies

The A_II_ AC oscillation frequency depends on its membrane potential; the more depolarized A_II_ ACs are, the higher is the oscillation frequency (Choi *et al*., 2014). This suggests that the A_II_ AC membrane potential in the various CSNB mouse models is more depolarized than in WT, with *Cav1*.*4*^*-/-*^ A_II_ ACs being the most depolarized and *mGluR6*^*-/-*^ the least. A_II_ ACs are ON-cells meaning that they depolarize with the reduction of glutamate release by the photoreceptors either via a direct ON-BC input or via a crossover inhibitory input from the OFF-BC pathway. Blocking all glutamate release from the photoreceptors, as is that case in the *Cav1*.*4*^*-/-*^ mice, would lead to a strong depolarization of the A_II_ ACs and thus a high oscillation frequency. Indeed, the oscillation frequency in the *Cav1*.*4*^*-/-*^ animals is the highest. In *Nyx*^*nob*^ animals the A_II_ ACs seem to be less depolarized, because the crossover inhibition from the OFF-BCs is still intact, which keep the A_II_ ACs somewhat more hyperpolarized compared to the *Cav1*.*4*^*-/-*^ mice. Our results would also suggest that the A_II_ ACs in *mGluR6*^*-/-*^ are less depolarize than they are in *Nyx*^*nob*^. We can only speculate why this may be so, but presumably the absence of the mGluR6 receptor affects the synapse differently than does the absence of the scaffolding protein nyctalopin. Unfortunately, the available literature regarding the exact resting membrane potentials of ON-BCs in the CSNB mouse models is contradictory (Tagawa *et al*., 1999; Ishii *et al*., 2009; Xu *et al*., 2012). However, following the arguments of Ishii et al., 2009, O’Conner et al., 2006 and Tagawa et al., 1999, it seems likely that ON-bipolar cells in the CSNB mouse models studied are constitutively depolarized.

#### 2) Why are *mGluR6*^*-/-*^ RGCs not oscillating spontaneously in the dark?

We propose that in the dark the *mGluR6*^*-/-*^ A_II_ ACs are resting at potentials just below that required to initiate spontaneous membrane potential oscillations. However, when stimulated with light they depolarize further and start to oscillate. This could explain why we and others (Takeuchi *et al*., 2018) find so few oscillating RGCs in the *mGluR6*^*-/-*^ retina compared with the other CSNB mouse models.

### 3) Why do RGCs and eyes in the *Cav1*.*4*^*-/-*^ mice oscillate under all stimulus conditions?

*Cav1*.*4*^*-/-*^ mice are blind and our results indicate their A_II_ ACs are strongly depolarized and oscillate with a degree of inherent synchronisation. This suggests that the A_II_ ACs coupling strength is higher in the *Cav1*.*4*^*-/-*^ retinas than in the other CSNB mouse models. The electrical coupling strength of neurons depends on the ratio of the gap-junction conductance and the input conductance of the coupled neurons (W. Kamermans, in preparation).

In *Cav1*.*4*^*-/-*^, crossover inhibition from the OFF pathway to the A_II_ ACs is also impaired and thus the associated inhibitory conductance will be closed. Accordingly, the input resistance of the A_II_ ACs in *Cav1*.*4*^*-/-*^ is likely to be higher than for the other CSNB mouse models. The reciprocal decrease in input conductance shifts the ratio towards the gap-junction conductance, increasing the electrical coupling strength between coupled A_II_ ACs. This increase in coupling between oscillating A_II_ ACs in *Cav1*.*4*^*-/-*^ enables the occurrence of an entrained oscillation state across the A_II_ AC network.

One prediction of the proposed mechanism is, that the oscillation frequency of the RGCs should depend on the light intensity. Indeed, the oscillation frequency in the *Nyx*^*nob*^ RGC increases with a higher light intensity and therefore the membrane potential of the A_II_ AC.

### Causal relation between RGC and eye movement oscillations

Consistent with previous reports (Michalakis *et al*., 2014), our results indicate *Cav1*.*4*^*-/-*^ mice are blind and as such their visual impairment is greater than is found in human patients with similar mutations. Since the photoreceptor output is absent with this mutation, it is interesting to compare results of the present study with those of the DNQX + D-AP5 experiments in *Nyx*^*nob*^ mice we described earlier (Winkelman *et al*., 2019). There we showed that applying this drug-cocktail to isolated retinas to block RGC inputs stops all RGC oscillations and injecting it into the eyes of intact mice stopped their eye movement oscillations. A result consistent with our retinal origin of nystagmus in CSNB hypothesis. However, this experiment does not per se discriminate between two possible conditions. The result could have occurred because blocking RGC inputs opened the oscillating OKR loop or because a retinal oscillator was silenced.

For the *Cav1*.*4*^*-/-*^ mice, their mutation will also open the OKR loop but now at the site of the photoreceptors, while leaving the RGC inputs intact. Hence, the output from oscillating A_II_ ACs is not blocked and still reaches the RGCs. The finding that these mice still have oscillating eye movements, shows that the oscillating A_II_ ACs are driving the nystagmus. Thus, although we cannot exclude some secondary modifications of the eye movement system downstream, these experiments confirm the retinal origin of the nystagmus in the CSNB mouse models and a causal relation between RGC and eye-movement oscillations.

### Differences in RGC and eye movement oscillation frequency

While there is a clear linear relationship between the RGC and eye-movement oscillations, the RGC oscillation frequencies are slightly higher than the frequencies for eye movements. This difference is most likely caused by the different physiological conditions of the in vitro and in vivo experiments. Many factors like pH, the composition of the external solution, temperature and potential retinal damage during the dissection may have influenced the RGC oscillations during the in vitro electrophysiological recordings.

### Mechanism potentially underlying other forms of nystagmus

The retinal mechanism we propose to cause nystagmus is most likely not only limited to CSNB since any condition that can depolarize A_II_ ACs beyond their normal working range will lead them to oscillate, possibly introducing nystagmus. Interestingly, a specific mutation in Munc 18-1 leads also to nystagmus (Li *et al*., 2020). Munc 18-1 is involved in the docking of synaptic vesicles to the membrane. It does so by binding with syntaxin. In the photoreceptor synaptic terminal Munc 18-1 binds to syntaxin 3B (Curtis *et al*., 2010). The mutation leading to nystagmus enhances the binding of Munc 18-1 with syntaxin 3B (Li *et al*., 2020). It is most likely that this mutation leads to reduced glutamate release by the photoreceptors resulting in depolarization of the A_II_ ACs, which in turn will start to oscillate and eventually lead to nystagmus. This example illustrates that it is essential to investigate whether novel genes implicated in nystagmus may have affected the A_II_ AC membrane potential and in that way, have induced nystagmus. This knowledge will allow to limit future therapeutic strategies to one common.

## Author Contributions

Conceptualization: M.B.H, M.H., B.H.W. and M.K.

Methodology: M.B.H, M.H., B.H.W.

Data curation: M.B.H, M.H., B.H.W. and M.K.

Formal analysis: M.B.H, B.H.W., M.H.

Funding acquisition: C.I.D.Z, M.K.

Investigation: M.B.H, M.H., B.H.W. and M.K.

Project administration: M.K.

Software: M.H., B.H.W. Supervision: C.I.D.Z, M. K.

writing—original draft preparation: M.B.H.

writing—review and editing: M.B.H, M.H., B.H.W., W.K., C.I.D.Z. and M.K.

All authors have read and agreed to the published version of the manuscript.

## Acknowledgements

We would like to thank Cynthia Geelen for genotyping the mice.

